# Inter-haplotype inversions and repeat expansion in the sexually deceptive orchid *Chiloglottis trapeziformis*

**DOI:** 10.64898/2026.05.25.727757

**Authors:** Zirui Zhang, Ashley Jones, Benjamin Schwessinger, Rod Peakall, Darren Wong

**Affiliations:** Research School of Biology, The Australian National University, Canberra, 2600, Australia; School of Agriculture, Food, and Wine, Waite Research Precinct, Adelaide University, Adelaide, SA 5064, Australia

**Keywords:** Genome evolution, Haplotype-resolved genome assembly, Structural variation, Chromosomal inversions, Transposable elements, Orchid

## Abstract

*Chiloglottis trapeziformis* is a sexually deceptive Australian orchid that provides a valuable system for studying orchid genome evolution, structural variation, and the molecular basis of specialized pollination. However, high-quality nuclear genome resources remain scarce for most orchids, particularly Australia’s diverse terrestrial lineages. To address this gap, we integrated PacBio HiFi, Oxford Nanopore ultra-long reads, and Hi-C chromatin-contact data to generate the first chromosome-scale, haplotype-resolved nuclear genome assembly for an Australian terrestrial orchid, *Chiloglottis trapeziformis*. Hi-C guided scaffolding resolved two haplotypes into 20 chromosomes each, consistent with the reported karyotype and genome size (2n=40, haplotype sizes of 1.58 Gb and 1.91 Gb). Genome completeness was high for both haplotypes, recovering 95.1% and 95.5% complete BUSCO genes for haplotype 1 and haplotype 2, respectively. De novo repeat annotation revealed a repeat-rich genome (85.79– 88.25% repetitive sequence), dominated by LTR retrotransposons. Evidence-guided annotation identified 16,287 and 16,548 protein-coding genes in Haplotype 1 and Haplotype 2, respectively. Phylogenetically informed comparisons placed *C. trapeziformis* as sister to *Anoectochilus roxburghii* among sampled Orchidoideae and showed broad gene-order conservation. Comparing the two haplotypes for structural variation, we identified large inter-haplotype inversions containing functionally annotated genes with detectable RNA expression, with focal examples further supported by local Hi-C contact patterns and breakpoint-level inspection. Inversion-overlapping genes did not show elevated dS relative to collinear background. This assembly and annotation resource provides a foundation for population and conservation genomics, structural and comparative analyses, and genome-enabled hypothesis testing of molecular traits underlying sexual deception in orchids.

**Significance statement:** Sexually deceptive orchids use remarkable chemical, visual, and tactile mimicry to attract specific pollinators, but the genome resources needed to understand how these complex traits evolved remain scarce. By generating a chromosome-scale, haplotype-resolved genome for *Chiloglottis trapeziformis*, we provide the first genomic framework for the large and unique Australasian tribe Diurideae. We show extensive structural variation between haplotypes in an otherwise highly collinear genome, with several large inversions potentially impacting expressed genes. The absence of elevated coding divergence in these regions highlights structural variation as a potentially underappreciated driver of orchid genome evolution. This resource fills a major gap for Australian orchids and provides a foundation for linking genome structure with orchid diversification, conservation, and the evolution of sexual deception.

## Introduction

With more than 25,000 species, Orchidaceae is among the most species rich families of flowering plants (Chase et al. 2015). The family is particularly well known for its contrasting epiphytic and terrestrial forms, extraordinary floral diversity and associated specialized pollination (Ackerman et al. 2023), dependence on mycorrhizal associations (Zhao et al. 2024), and repeated evolution of crassulacean acid metabolism (CAM) photosynthesis (Gamisch et al. 2021). These adaptive traits are predicted to be associated with large, and structurally complex orchid genomes (Trávníček et al. 2019).

The publication of the first orchid reference genome, *Phalaenopsis equestris* (Cai et al. 2015), marked a turning point in our ability to test this prediction. Substantial genomic progress has since been made, particularly in the large epiphytic subfamily (∼16 tribes, ∼600 genera, ∼20,500 sp), Epidendroideae, where genomes span 10 genera, and a pangenome exists for the large genus *Dendrobium* (Li et al. 2025). Nonetheless, whole-genome resources for the five other orchid subfamilies with mostly terrestrial species remain sparse (Zhang et al. 2022). In the next in size subfamily, the Orchidoideae (∼4 tribes, ∼190 genera, ∼4300 sp), genomes are only available for one member of the Cranichideae (∼1500 sp) (*Anoectochilus*, Xu et al. 2025), and four members of the Orchideae (∼1800 sp): a *Dactylorhiza* (Wolfe et al. 2023) an *Ophrys* (Russo et al. 2024), and two *Platanthera* (Li et al. 2022). Presently, there are no published reference nuclear genomes representing the distinctive Australasian tribe the Diurideae (∼950 sp) or more than 200 other southern hemisphere representatives of the subfamily.

The Australasian terrestrial genus *Chiloglottis* (Orchidoideae: Diurideae, ∼30 sp) offers a powerful first case study to address this genomic resource gap. Like several other sexually deceptive Australian orchid genera, *Chiloglottis* orchids attract specific male thynnine wasps as pollinators by mimicking the sex pheromones, appearance, and tactile cues of the female wasps (Wong et al. 2019). With detailed studies spanning pollination, chemistry, biosynthesis, genetics and phylogenomics of the group (Wong et al.2022a; 2022b), it has become a model lineage for studying the evolution of sexual deception and other key traits. Here, using PacBio HiFi reads, Oxford Nanopore ultra-long reads, and Hi-C chromosome conformation data, we report a high-quality chromosome-scale genome assembly for *Chiloglottis trapeziformis* (Fig. 1A). The resulting assembly comprises two haplotypes, each resolved into 20 chromosomes, and is accompanied by repeat annotation, gene prediction, and functional annotation. This genome provides the essential foundation for future molecular, genomic, and comparative evolutionary studies of *Chiloglottis* and other southern hemisphere terrestrial orchids.

**Figure 1.**
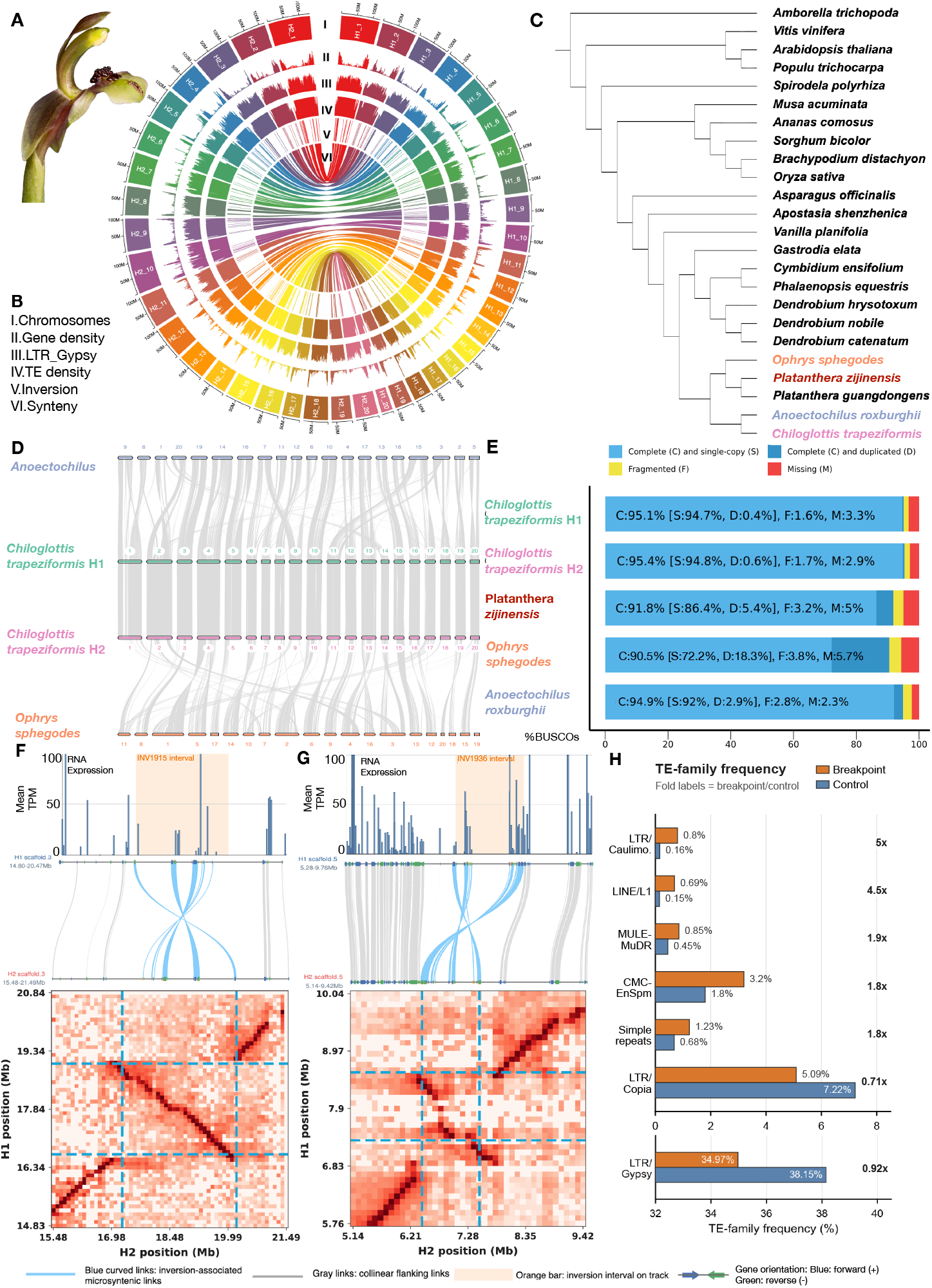
Chromosome-scale genome features, comparative genomics, and representative inversions in *Chiloglottis trapeziformis*. (A) Floral morphology of *C. trapeziformis*. (B) Circos plot depicting the distribution of i. chromosome, ii. gene density, iii. *Gypsy* long terminal repeats (LTR), iv. Transposable element (TE) density, v. inversion occurrence, and vi. synteny across assembled *C. trapeziformis* haplotypes. (C) Species tree depicting phylogenetic placement of *C. trapeziformis* among representative angiosperms, with emphasis on orchid lineages (all nodes have bootstrap support of 100). (D) Synteny among *Anoectochilus roxburghii, C. trapeziformis* haplotypes 1 and 2 (H1, H2), and *Ophrys sphegodes*, highlighting collinear flanking regions and inversion-associated links. (E) BUSCO assessment of assembly completeness for the *C. trapeziformis* haplotypes and selected orchid genomes. (F, G) Two representative high-confidence inversions (INV1915 and INV1936). In each panel, the upper track shows mean RNA-seq expression across annotated genes on the interval, with the shaded orange region marking the inversion interval. The middle track shows local microsynteny between the corresponding H1 and H2 regions, with blue links indicating inversion-associated gene pairs and grey links indicating collinear flanking regions; gene arrows show orientation. The lower heatmap shows the local Hi-C contact pattern across the H1 and H2 coordinates, with dashed blue boxes highlighting the inferred inversion boundaries. (H) Comparison of TE-family frequencies between inversion breakpoint windows and randomly sampled control regions, showing that breakpoint composition differs from the genomic background.

## Results and Discussion

### Hybrid sequencing strategy for a chromosome-scale, haplotype-resolved assembly

Resolving large, repeat-rich and heterozygous plant genomes requires chromosome-scale, and ideally haplotype-resolved, assemblies because fragmented or collapsed assemblies can obscure annotation, variant detection, structural variation and repeat dynamics (Formenti et al. 2022). This is particularly relevant for orchids, which show exceptional genome-size variation, high repeat loads and substantial heterozygosity (Trávníček et al. 2019; Zhang et al. 2022). We therefore generated a chromosome-scale, haplotype-resolved genome for *Chiloglottis trapeziformis* by integrating PacBio HiFi, Oxford Nanopore ultra-long reads and Hi-C data.

HiFi sequencing produced 79.97 Gb of accurate long reads (read N50 = 19.4 kb; median quality Q25), providing approximately 40× coverage per 2 Gb assumed haplotype. ONT sequencing generated 200.34 Gb of raw sequence data; post-filtering, 159.96 Gb of reads remained (read N50 = 49.8 kb; median Q17.4), yielding ∼80× coverage per haplotype. Finally, Hi-C sequencing yielded 141.14 Gb of 2×150 bp paired-end data (138.38 Gb post-filtering; median Q28.7), providing long-range linkage data essential for chromosomal scaffolding (Table S1).

### A highly contiguous, gene-complete reference resource for *C. trapeziformis*

Genome assembly with hifiasm (UL) yielded long contiguous contigs with robust gene-space completeness across both haplotypes: haplotype 1 (H1; 443 contigs, 1.69 Gb, N50 = 32.7 Mb) and haplotype 2 (H2; 281 contigs, 2.08 Gb, N50 = 42.8 Mb; Table S2). The largest contigs reached 66.6 Mb (H1) and 106.2 Mb (H2), demonstrating long-range continuity. BUSCO assessment (embryophyta_odb12) confirmed high gene-space completeness with 94.61% and 95.29% complete BUSCOs, respectively. Coverage of HiFi reads across the chromosomes was consistently high, being 22.18× and 21.44× for haplotype 1 and 2 respectively (Fig. S1; Table S3). Following contamination screening, most retained contigs with confident taxonomic assignments matched known orchid sequences, including 69.23% in H1 and 62.67% in H2. Hi-C scaffolding resolved both assemblies into 20 chromosomes (Fig. 1B; Fig. S2), consistent with the reported *Chiloglottis* chromosome count (2n=40) (Peakall and James 1989; Dawson et al. 2007). Most assembly sequence was anchored into chromosome-scale scaffolds for both haplotypes: 93.73% in H1 (1.58 Gb; scaffold N50 = 86.4 Mb) and 91.88% in H2 (1.91 Gb; scaffold N50 = 100.6 Mb) (Table 1).

**Table 1.** Summary of the *Chiloglottis trapeziformis* genome assembly and annotation.

| Statistic | Haplotype 1 | Haplotype 2 |
| --- | --- | --- |
| **Chromosome-scale scaffolding** |  |  |
| Chromosome scaffolds | 20 | 20 |
| Total length (bp) | 1,581,935,574 | 1,908,350,318 |
| Largest contigs (bp) | 122,431,139 | 134,196,054 |
| N50 (bp) | 86,375,386 | 100,600,513 |
| **Repeat composition** |  |  |
| Repetitive sequence (%) | 85.79 | 88.25 |
| Transposable elements (bp; %) | 746,252,173 (47.18) | 1,072,643,715 (56.21) |
| Retrotransposons (Class I; %) | 44.55 | 54.21 |
| Gypsy/DIRS1 (%) | 36.97 | 38.77 |
| DNA transposons (Class II; %) | 2.63 | 2.00 |

| Gene Annotation |  |  |
| --- | --- | --- |
| Protein-coding genes | 16,287 | 16,548 |
| Total transcripts | 22,081 | 22,387 |
| BUSCO complete (C = S + D; %) | 95.06 | 95.46 |
| BUSCO complete single-copy (S; %) | 94.67% | 94.82% |
| BUSCO complete duplicated (D; %) | 0.39% | 0.64% |
| BUSCO non-complete (F + M; %) | 4.94% | 4.54% |

### *De novo* repeat annotation revealed an exceptionally repeat-rich *C. trapeziformis* genome

Consistent with the high repeat content reported for many orchid genomes (Zhang et al. 2022), repetitive regions dominated across the *C. trapeziformis* genome (H1, 85.79%; H2, 88.25%; Table 1; Table S4). These values fall within the upper range of orchid repeat abundance, comparable to the *Cymbidium goeringii* genome (88.87%; Chung et al. 2022) and exceeding other orchids. Transposable elements (TEs) accounted for a large share of the repetitive DNA (H1, 47.18%, 746 Mb; H2, 56.21%, 1,073 Mb), with long terminal repeat (LTR) elements contributing most TE sequences, primarily from the Gypsy/DIRS1 subclass (Table 1). Nonetheless, a substantial fraction of the repetitive DNA of *C. trapeziformis* remained unclassified (31–38%). High proportions of unclassified repeats have also been reported in other orchid genomes (e.g. *Phalaenopsis equestris*, 21.99%, Cai et al. 2015; *Gongora gibba*, 30.88%, Guizar Amador et al. 2024). Thus a substantial fraction of orchid repetitive DNA remains difficult to assign to established categories, suggesting the presence of lineage-specific repeat families or limited representation of specific orchid repeats in curated databases.

### Evidence-guided annotation reveals a highly complete gene landscape across haplotypes

Integrating *ab initio* prediction, protein homology, and earlier RNA-seq evidence from multiple floral tissues and developmental stages, we identified comprehensive protein-coding gene sets for both haplotypes. Gene prediction recovered similar numbers of gene models in H1 and H2, with high BUSCO completeness (H1, 16,287 genes, 95.06% complete BUSCOs; H2, 16,548 genes, 95.46% complete BUSCOs; Table S5). MapMan4 assigned functional categories to 77.92% of predicted genes, consistent with other orchid genomes in which lineage-specific sequences and database biases toward model species can limit annotation (Fig. S3) (Guizar Amador et al. 2024). These gene and functional annotations provide a foundation for downstream pathway-level and comparative analyses in Australian sexually deceptive orchids.

### Phylogenetically informed assessment of genome quality and synteny

The phylogenetic placement of *C. trapeziformis* within Orchidoideae and as sister to *Anoectochilus roxburghii* among sampled taxa matched expectations (Peakall et al. 2021; Wong and Peakall 2022; O’Donnell et al. 2025; Fig. 1C; Fig. S4). Gene-based synteny comparisons further supported the structural quality and comparative utility of the assembly: extensive collinearity was observed between the two *C. trapeziformis* haplotypes and between each haplotype and other Orchidoideae genomes, with the strongest syntenic correspondence to *A. roxburghii* and more fragmented collinearity against the more divergent *Ophrys* genome (Fig.1D; Table S6). Comparative BUSCO assessment showed that both *C. trapeziformis* haplotypes had high gene-space completeness relative to other sampled orchid genomes (Fig. 1E).

### Inversions show TE-associated breakpoint signatures without elevated synonymous divergence

Despite extensive large-scale collinearity between the two haplotypes within *C. trapeziformis* and with related species, synteny analysis revealed several local inter-haplotype inversions (Fig. 1F). We identified a prioritized set of seven inversions whose genic regions had detectable floral RNA-seq expression (Table S7). Of these, INV1915 and INV1936 offer representative examples for further investigation. INV1915 represents a clear structural inversion whose genic regions are associated with ribosome biogenesis, chromatin regulation, lipid metabolism, and mitochondrial cofactor assembly, but without obvious breakpoint disruption of any annotated gene. By contrast, INV1936 illustrates a more complex breakpoint architecture whose associated genes include protein biosynthesis and turnover, membrane transport, and signal transduction. Closer inspection of INV1936 showed that the inversion boundary intersects an orthologous gene pair (g13041 in H1 and g13243 in H2) in both haplotypes, 536 bp from the nearest exon boundary (Fig. 1G; Fig. S5). Although the boundary-separated terminal segment spans 1,359 bp and includes 441 bp of coding sequence, the mirrored breakpoint configuration suggests that neither haplotype carries a uniquely truncated gene. Together, gene annotations and RNA-seq support are more consistent with possible local regulatory or splicing-related effects than with direct coding-sequence loss.

The evolutionary consequences of inversions vary substantially across plants. At a broader orchid scale, genus-wide analyses of *Dendrobium* identified both shared and lineage-specific inversions, highlighting structural dynamics among closely related taxa (Li et al. 2025). Shifting to other monocots, inverted regions in *Curcuma* have been associated with relaxed genetic constraints, elevated substitution rates, and disruption of tandemly duplicated TPS genes involved in terpenoid biosynthesis (Liao et al. 2024). Conversely, in the eudicot daisy *Gorteria diffusa*, large chromosomal inversions drive phenotype stabilization by suppressing recombination in hybrid zones, thereby preserving tightly integrated floral traits crucial for sexual deception and reproductive fitness (Kellenberger et al. 2026). In contrast to these highly dynamic systems, our coding-divergence analyses detected no elevation of synonymous divergence (dS) for inversion-associated genes relative to collinear background regions (Table S8), consistent with models in which inversions primarily influence evolution through recombination suppression and altered linkage rather than immediate coding divergence (Wellenreuther and Bernatchez 2018; Huang and Rieseberg 2020).

Interestingly, breakpoint regions in *C. trapeziformis* showed family-specific shifts in transposable-element composition rather than generalized repeat accumulation. Several TE families were enriched relative to the genomic background, including LTR/Caulimovirus (5.0-fold within ±5 kb), LINE/L1 (4.5-fold), MULE-MuDR (1.9-fold), and CMC-EnSpm (1.8-fold), whereas the genome-dominant LTR/Gypsy and LTR/Copia families were depleted (0.92-fold and 0.71-fold, respectively) (Fig. 1H; Table S9). Because enriched categories contribute only a minor fraction of total repeat bp, these patterns likely reflect compositional shifts rather than global repeat enrichment, consistent with mechanisms involving transposon-mediated alternative transposition, repeat-mediated rearrangement, or ectopic recombination (Sharma and Peterson 2023; Zhou et al. 2023). In the future, individual-level, haplotype-resolved transcriptomic analyses matched across tissues and developmental stages will be required to test whether focal inversions are associated with cis-regulatory divergence, despite limited coding-sequence differentiation.

In conclusion, we generated a high-quality haplotype-resolved reference genome for the sexually deceptive orchid *C. trapeziformis*. Its placement within an underrepresented Australasian terrestrial orchid lineage, together with the identification of large inter-haplotype inversions, highlights the value of this assembly for future studies of orchid genome evolution, structural variation, and conservation genomics. Beyond our focal system, this genomic resource provides the necessary foundation for future investigations into the evolution of sexual deception by establishing a comparative framework to resolve how such complex pollination strategies arise in orchids (Russo et al. 2024) and across broader plant lineages where sexual mimicry has independently evolved (Kellenberger et al. 2026).

## Materials and Methods

### Sampling, DNA extraction, and sequencing

Fresh young leaves of *Chiloglottis trapeziformis* were collected from a single clone of plants cultivated at the Royal Botanic Gardens Victoria (Melbourne, Australia) and snap-frozen at −80 °C. High-molecular-weight DNA was extracted using a magnetic bead-based protocol (Jones et al. 2021). Prior to sequencing, the DNA size-selected for fragments ≥20 kb using a BluPippin (Sage Science). PacBio HiFi libraries were prepared with the SMRTbell Prep Kit v3.0 and sequenced on a PacBio Revio platform. Oxford Nanopore libraries were prepared using the Ligation Sequencing Kit v14 and sequenced on a PromethION P24 (R10.4.1 flow cells). Hi-C libraries were prepared using the Phase Genomics Proximo Hi-C Plant Kit v4 and sequenced on an Illumina NovaSeq 6000 (2 × 150 bp). Sequencing reads were processed using standard quality-control procedures (details in Supplementary Methods).

### Genome profiling and haplotype-resolved assembly

Genome ploidy and heterozygosity inferred using Smudgeplot (Fig. S6). The homozygous coverage peak was set to 48× from the HiFi k-mer profile and used as the hifiasm --hom-cov parameter. A haplotype-resolved diploid assembly was generated using hifiasm v0.19.8 (Cheng et al. 2024), integrating PacBio HiFi, Oxford Nanopore ultra-long, and Hi-C reads. Assembly quality was assessed using Bandage (Wick et al. 2015) and BUSCO via compleasm (embryophyta_odb12) (Seppey et al. 2019), with the optimal assembly selected based on contiguity, gene completeness, and concordance with k-mer–based genome size estimates. Contigs were screened against NCBI nt to remove non-plant sequences (details in Supplementary Methods). Filtered assemblies were scaffolded with Hi-C data using Bowtie2 v2.5.4 (Langmead and Salzberg 2012), HiC-Pro v3.1.0 (Servant et al. 2015), and YaHS v1.2a (Zhou et al. 2022). Hi-C contact maps were inspected and manually curated with JuicerTools v3.0 and Juicebox v2.20 (Durand et al. 2016). Each haplotype was resolved into 20 pseudochromosomes, consistent with the diploid chromosome count across *Chiloglottis* (Peakall and James 1989; Dawson et al. 2007). (details in Supplementary Methods).

### Assembly validation and genome annotation

Genome and gene completeness of each haplotype was assessed using BUSCO via compleasm v0.2.7 (embryophyta_odb12). Inter-haplotype structural concordance was evaluated using D-GENIES for whole-genome alignment visualisation (Cabanettes and Klopp 2018). PacBio HiFi reads were mapped back to scaffolded assemblies using minimap2 (Li 2021), and per-scaffold coverage was quantified with BamToCov v2.7.0 (Birolo and Telatin 2022) to assess coverage uniformity and scaffold integrity. Haplotype-specific repeat libraries were generated using RepeatModeler v2.0.5 (Flynn et al. 2020) and annotated with RepeatMasker v4.1.5 (Chen 2004). Protein-coding genes were predicted using BRAKER3 v3.0.8 in evidence-guided mode (Gabriel et al. 2024)and functionally annotated with MapMan4 BINs (Schwacke et al. 2019) (details in Supplementary Methods).

### Phylogenomic and comparative genomic analyses

Phylogenomic species-tree inference was performed using OrthoFinder v2.5.5 (Emms and Kelly 2019) with minor modification. The orchid genome resource survey and plant genomes used for phylogenomic and comparative analyses are summarized in Tables S10 and S11. Comparative analyses leveraged sequenced Orchidoideae relatives (*Anoectochilus roxburghii, Ophrys sphegodes*, and *Platanthera zijinensis*). Genome completeness and functional annotation were re-assessed for each of these genomes as previously described for *C. trapeziformis* to ensure meaningful comparisons. Synteny relative to *C. trapeziformis* haplotypes was evaluated using the MCScan (Tang et al. 2024) (details in Supplementary Methods).

### Inter-haplotype inversion detection and evidence-based prioritization

Inter-haplotype whole-genome alignments were generated with minimap2 v2.26, and inversions were inferred with SyRI v1.6 (Goel et al. 2019). Hi-C reads were mapped to a joint H1+H2 reference and processed with Juicer v1.5 (Durand et al. 2016). Expression and functional evidence were integrated to prioritise high-confidence inversions and visualised using IGV (Thorvaldsdóttir et al. 2013). (details in Supplementary Methods).

## Acknowledgments

We acknowledge Australia’s First Nations Peoples as the Traditional Custodians of the land and waters on which this work was conducted, recognizing their historical and ongoing stewardship of its genetic resources, and pay our respects to all Elders past and present. We would like to thank the Biomolecular Resource Facility at the John Curtin School of Medical Research, ANU in Canberra, Australia, where sequencing was conducted.

The authors acknowledge the provision of computing and data resources provided by the Australian BioCommons Leadership Share (ABLeS) program. This program is co-funded by Bioplatforms Australia, enabled by the Commonwealth Government National Collaborative Research Infrastructure Strategy (NCRIS), the National Computational Infrastructure and Pawsey Supercomputing Research Centre (Gustafsson et al. 2023).

We are indebted to Dr Noushka Reiter (Royal Botanic Gardens Victoria) for access to propagated samples of *Chiloglottis trapeziformis* that were known to be of a single clonal origin, and to Monica Ruibal for sampling assistance.

## Data Availability

The *C. trapeziformis* haplotype-resolved genome assemblies have been submitted to NCBI as separate haplotype assemblies from the same BioSample (SAMN59625637). Haplotype 1 is associated with BioProject PRJNA1463801, and haplotype 2 is associated with BioProject PRJNA1468196. Final GenBank assembly accessions are pending NCBI processing and will be updated upon release. Additional supporting files, including genome annotation files, are available from Zenodo at https://doi.org/10.5281/zenodo.20321953.

