## supplementary methods for "Inter-haplotype inversions and repeat expansion in the sexually deceptive orchid *Chiloglottis trapeziformis*"

**Sampling, DNA extraction, and sequencing**

### Fresh young leaves of Chiloglottis trapeziformis were collected from plants cultivated at the Royal Botanic Gardens Victoria (Melbourne, Australia) and snap-frozen at −80°C. High-molecular-weight genomic DNA was extracted using a magnetic bead-based protocol (Jones et al., 2021). DNA integrity was assessed with a Femto Pulse (Agilent, Genomic DNA 165 kb kit), and fragments ≥20 kb were size-selected using BluePippin (Sage Science).

PacBio HiFi libraries were prepared with the SMRTbell Prep Kit 3.0 and sequenced on a Revio platform (25M SMRT Cell). Oxford Nanopore libraries were prepared using the Ligation Sequencing Kit V14 (SQK-LSK114) and sequenced on a PromethION P24 with two FLO-PRO114M R10.4.1 flow cells. Flow cells were washed, re-primed and reloaded as output declined. Raw ONT signal (POD5) was basecalled to FASTQ using Dorado v0.5.3 with the super-accurate model dna_r10.4.1_e8.2_400bps_sup@v4.3.0. Terminal 200 bp were trimmed from each end using Chopper v0.7.0, and reads were filtered to retain sequences ≥20 kb with mean quality ≥Q7. Read-length and quality distributions were summarised using NanoPlot2 v1.42.0.

For chromosome conformation capture, a Hi-C library was prepared using the Phase Genomics Proximo Hi-C Plant Kit v4 (KT3040B) and sequenced on an Illumina NovaSeq 6000 (2×150 bp). Hi-C reads were adapter-trimmed and quality-filtered using TrimGalore v0.6.10 (parameters: –quality 30, –length 100), retaining read pairs ≥100 bp after trimming.

**Genome profiling and haplotype-resolved assembly**

Genome size and *k*-mer abundance were estimated from filtered PacBio HiFi reads using KmerGenie v1.7051. Ploidy and heterozygosity were inferred using Smudgeplot v0.2.5. Based on *k*-mer spectra, the homozygous coverage peak was set to 48× for downstream assembly.

A haplotype-resolved diploid de novo assembly was generated using hifiasm v0.19.8 (Cheng et al. 2024). Filtered PacBio HiFi reads, filtered ONT ultra-long reads (option --ul) and filtered Hi-C read pairs (--h1 --h2) were used as input. Assembly graphs were inspected using Bandage v0.9.0 (Wick et al. 2015). Gene-space completeness was evaluated using BUSCO via compleasm v0.2.7 (embryophyta_odb12) (Seppey et al. 2019). When multiple assemblies were produced under different filtering conditions, candidate contig sets were compared based on contiguity (N50), BUSCO completeness (emphasising single-copy genes with low duplication) and concordance with the k-mer–based genome-size estimate, selecting the best assembly for scaffolding.

**Contamination screening**

To screen for non-plant contamination, contigs from each haplotype were queried using BLASTn v2.14.0 against the NCBI nucleotide (nt) database (downloaded 10 April 2024). Contigs were split into 1 Mb windows prior to BLAST queries. A contig was flagged as non-plant when the majority of high-confidence BLAST hits across windows were assigned to non-Viridiplantae taxa (e-value ≤1e−20; alignment length ≥500 bp; percent identity ≥80%) and was removed. Contigs with predominantly Viridiplantae hits or lacking high-confidence hits were retained.

### **Hi‑C–based chromosome-scale scaffolding**

Filtered assemblies were scaffolded using Hi-C data. Hi-C reads were aligned using Bowtie2 v2.5.4 (Langmead and Salzberg 2012), then processed with HiC-Pro v3.1.0 (Servant et al. 2015) to identify valid interaction pairs. Scaffolding was carried out with YaHS v1.2a (Zhou et al. 2022). Given the high continuity of long-read contigs, error correction during scaffolding was disabled to avoid over-fragmentation, and only high-confidence Hi-C alignments (MAPQ >20) were used. Hi-C contact maps were inspected with JuicerTools (v3.0) (Durand et al. 2016), and manual corrections were performed using Juicebox v2.20 when needed. After curation, each haplotype assembly was resolved into 20 pseudochromosomes, matching the diploid karyotype (2n = 40) reported for *Chiloglottis* (Peakall and James 1989; Dawson et al. 2007).

**Assembly quality assessment and validation**

Genome completeness for each haplotype was reassessed using BUSCO via compleasm v0.2.7 (embryophyta_odb12). Inter-haplotype structural concordance and potential large-scale discrepancies (e.g., inversions or translocations) were evaluated using D-GENIES (Cabanettes and Klopp 2018) for whole-genome alignment visualisation. To assess read-level support and coverage uniformity across pseudochromosomes, PacBio HiFi reads were mapped back to scaffolded assemblies using minimap2 (Li 2021). Coverage per scaffold was quantified using BamToCov v2.7.0 (Birolo and Telatin 2022). Consistent read depth across anchored pseudochromosomes and absence of extended zero-coverage regions were interpreted as evidence for scaffold integrity.

**Repeat annotation, gene prediction, and biological pathway annotation**

To support repeat-aware gene prediction, haplotype-specific *de novo* repeat libraries were generated using RepeatModeler v2.0.5 (Flynn et al. 2020). Repetitive sequences and transposable elements were annotated using RepeatMasker v4.1.5 (Chen 2004) with soft-masking enabled.

Protein-coding genes were annotated on soft-masked haplotype assemblies using BRAKER3 v3.0.8 in evidence-guided mode (Gabriel et al. 2024). Transcript evidence comprised 56 *Chiloglottis* RNA-seq datasets (NCBI SRA study accession: SRP109328 & SRP281918) spanning key floral developmental stages, chiloglottone (UV-B)-inducing treatments, and leaf tissues (Wong et al. 2017; Peakall et al. 2021). RNA-seq reads were adapter-trimmed and quality-filtered using fastp v0.20.0 (Chen 2023) prior to use. Protein homology evidence was provided using proteomes from 10 published orchid genomes (Tables S10 and S11), selected to improve annotation sensitivity within Orchidaceae and reduce spurious models driven by distant homology. Gene set completeness was assessed using BUSCO in gene mode via compleasm v0.2.6 (embryophyta_odb12). Functional annotation was performed using MapMan4 v6.0 BIN categories using Mercator webtool with default parameters (Schwacke et al. 2019).

**Species tree inference**

Phylogenomic analysis was performed as previously described with modifications (Wong and Peakall 2022). Briefly, orthogroups were inferred with OrthoFinder v2.5.5 (Emms and Kelly 2019) using the multiple-sequence-alignment workflow (-M msa; -I 1.3, -t 40, -a 8) on predicted proteins from *C. trapeziformis*, 12 additional orchid genomes, and 11 representative angiosperms. Species tree inference was based on a custom near-single-copy orthogroup set extracted from the OrthoFinder gene-count matrix. Near-single-copy orthogroups were defined as those with >=80% taxon occupancy, no more than two copies in any species, and no more than four multicopy species per orthogroup. This yielded 354 near-single-copy orthogroups and a concatenated supermatrix of 192,895 amino acid sites. Maximum-likelihood species tree inference was performed in IQ-TREE v2.3.5 (Minh et al. 2020) (-m MFP+MERGE, --B 1000,--alrt 1000, --bnni). Tree visualization was performed with iTOL v4 (Letunic and Bork 2019).

**Comparative genomics within Orchidoideae**

Based on the phylogenetic placement of *C. trapeziformis* within Orchidoideae, we selected the closest available sequenced relatives in our dataset for comparative evaluation, including *Anoectochilus roxburghii, Ophrys sphegodes*, and *Platanthera zijinensis*. We then compared genome completeness (BUSCO) via compleasm v0.2.7 (embryophyta_odb12), functional annotation using Mercator webtool with default parameters (Schwacke et al. 2019), and gene-based synteny patterns relative to *C. trapeziformis* haplotypes. Synteny was assessed using the MCScan workflow implemented in JCVI utilities v1.4.21 (Tang et al. 2024). Pairwise similarity searches were performed using LAST v1550 (Kiełbasa et al. 2011).

**Inter-haplotype inversion detection and evidence-based prioritization**

To provide independent physical support for each inversion, we constructed a joint H1+H2 reference by concatenating the top 20 scaffolds from each haplotype assembly with haplotype-prefixed chromosome identifiers. Hi-C reads were mapped to this joint reference using BWA-MEM v0.7.17 (-SP5M), followed by chimeric read classification, fragment assignment and PCR duplicate removal using Juicer v1.5 core scripts (Durand et al. 2016), with a positional wobble tolerance of 4 bp for duplicate detection. Deduplicated read pairs were converted into a multi-resolution .hic contact matrix using juicer_tools pre v1.9.9 with no MAPQ filtering (MAPQ >= 0). For each candidate inversion, inter-haplotype contact frequencies within local windows were extracted using juicer_tools dump observed at 100 kb resolution with Knight-Ruiz (KR) matrix balancing normalization (Knight and Ruiz 2013), with Vanilla Coverage Square Root (VC_SQRT) as a fallback. Contact values were log-transformed (log(1 + x)) and visualized as heatmaps. Inversion boundaries were overlaid as dashed lines; in a correctly identified inversion, inter-haplotype contacts form an anti-diagonal pattern within the inverted region flanked by diagonal signal in collinear regions.

### Genes overlapping each inversion were identified using the H1 BRAKER gene annotation. RNA-seq reads from the datasets used for genome annotation were aligned to the H1 reference with minimap2 (-ax splice --secondary=no) and quantified per gene using featureCounts with the BRAKER GTF. To reduce developmental and tissue heterogeneity, inversion-expression analyses were restricted to a stage- and tissue-specific floral subset comprising three sunflower-stage labellum libraries. Raw read counts from this subset were converted to transcripts per million (TPM) by dividing each gene's count by its length in kilobases to obtain reads per kilobase (RPK), then dividing by the per-sample sum of RPK scaled to millions. Expression values were mapped to inversion-overlapping genes and summarized as per-inversion mean, median, and sum statistics.

### The same sunflower-stage labellum subset was used for local expression visualization of the focal inversion cases INV1915 and INV1936. For breakpoint-level inspection of INV1936, all available RNA-seq libraries were additionally merged to maximize local junction and read-coverage support; these merged data were used only for qualitative transcript-structure inspection rather than formal expression quantification. MapMan4 functional annotations of H1 and H2 gene models were used to assign putative biological functions to inversion-overlapping genes.

Hi-C contact map support, gene overlap counts, RNA expression summaries and MapMan4 functional annotations were merged into a unified multi-sheet evidence table. The merged evidence table was used to identify a high-confidence subset of seven inversions by integrating local Hi-C context, genic context, detectable expression of overlapping genes, and functional annotation. Two of these inversions, INV1915 and INV1936, were selected as representative focal examples for visualization in the main text because they showed clear structural support and contrasting breakpoint contexts.

**Codon-based divergence of inversion-associated genes**

To quantify sequence divergence between the two Chiloglottis haplotypes, we analyzed anchor-defined one-to-one orthologous gene pairs between the H1 and H2 assemblies using the primary transcript models (.t1). Syntenic H1-H2 anchor pairs were obtained from the H1t1.H2t1.anchors file, and only pairs unique on both haplotypes were retained. CDS sequences for each anchored gene pair were extracted from the H1 and H2 CDS files, converted to uppercase, non-ACGTN characters were replaced with N, and terminal stop codons were removed when present. CDSs were translated with the standard genetic code; when peptide lengths differed, amino-acid sequences were globally aligned and the alignment was back-translated to a codon alignment. Codons containing gaps in either sequence were excluded from identity and distance calculations. Synonymous (dS) and nonsynonymous (dN) substitution rates were then estimated from codon alignments using the Nei-Gojobori (NG86) method implemented in Biopython, and dN/dS was calculated where dS was finite. Gene pairs containing internal stop codons were excluded from dN/dS estimation.

For the inversion-overlapping genes, we used the SyRI inversion set and identified H1 genes fully contained within each inversion interval. These inversion-interior genes were intersected with the precomputed one-to-one anchor divergence table for downstream analysis. As controls, we used non-inversion collinear anchor pairs, and for matched comparisons each inversion gene was paired without replacement to a control gene from the same chromosome with the closest CDS length. Distributions of dS, dN, dN/dS, and sequence-distance metrics were compared using two-sided Mann-Whitney U tests, and paired inversion-versus-control comparisons were additionally assessed with two-sided Wilcoxon signed-rank tests.

**Repeat and transposable element analysis at inversion breakpoints.**

To characterise the repeat landscape at inversion breakpoints and test whether specific TE families are associated with breakpoint regions, we compared the repeat composition of breakpoint-flanking windows with size-matched random genomic controls. For each of the 171 inversions, two breakpoint positions were defined on the H1 reference (the left and right boundaries), yielding 342 breakpoint loci. RepeatMasker annotations (H1 genome; RepeatMasker v4.1.5, rmblastn v2.13.0) were parsed and indexed by chromosome. For each breakpoint, a window of size w was centred on the breakpoint position (clamped to chromosome boundaries), and all overlapping RepeatMasker entries were intersected with the window to compute (i) the total repeat fraction (merged repeat base pairs divided by window length) and (ii) base-pair coverage attributed to each TE class (e.g. DNA, LINE, LTR) and family (e.g. LTR/Gypsy, DNA/CMC-EnSpm). For comparison, 100 random control windows of the same size per breakpoint (34,200 total) were sampled from the same chromosomes, excluding inversion-interior regions, using a fixed random seed. Overall repeat density was compared between the 342 breakpoint windows and 34,200 control windows using two-sided Mann–Whitney U tests. TE class- and family-level composition was summarised as the percentage of total window base pairs attributed to each category, and fold enrichment was calculated as the ratio of breakpoint percentage to control percentage. Analyses were performed at three window sizes (±5 kb, ±10 kb, and ±20 kb) to assess robustness of the enrichment signal.
